# Type-2 diabetes mellitus enhances *Klebsiella pneumoniae* pathogenesis

**DOI:** 10.1101/2024.05.31.596766

**Authors:** Katlyn Todd, Krista Gunter, James M. Bowen, Caitlyn L. Holmes, Natasha L. Tilston-Lunel, Jay Vornhagen

**Author notes:** Corresponding author: Jay Vornhagen.

## Abstract

*Klebsiella pneumoniae* is an opportunistic pathogen and an important cause of pneumonia, bacteremia, and urinary tract infection. *K. pneumoniae* infections are historically associated with diabetes mellitus. There is a fundamental gap in our understanding of how diabetes mellitus, specifically type 2 diabetes, influences *K. pneumoniae* pathogenesis. *K. pneumoniae* pathogenesis is a multifactorial process that often begins with gut colonization, followed by an escape from the gut to peripheral sites, leading to host damage and infection. We hypothesized that type 2 diabetes enhances *K. pneumoniae* pathogenesis. To test this, we used well-established mouse models of *K. pneumoniae* colonization and lung infection in conjunction with a mouse model of spontaneous type 2 diabetes mellitus (T2DM). We show that T2DM enhances susceptibility to both *K. pneumoniae* colonization and infection. The enhancement of gut colonization is dependent on T2DM-induced modulation of the gut microbiota community structure. In contrast, lung infection is exacerbated by the increased availability of amino acids in the lung, which is associated with higher levels of vascular endothelial growth factor. These data lay the foundation for mechanistic interrogation of the relationship between *K. pneumoniae* pathogenesis and type 2 diabetes mellitus, and explicitly establish T2DM as a risk factor for *K. pneumoniae* disease.

## Introduction

Type 2 diabetes mellitus (T2DM), which is characterized by hyperglycemia due to insulin resistance, increases the risk of infection and mortality due to infection [1-7] and is rising in prevalence [8]. Diabetic individuals are more susceptible than non-diabetics to infection due to immune dysregulation (reviewed in [9, 10]) and are more susceptible to colonization by opportunistic pathogens [11, 12]. T2DM significantly alters biological processes necessary for controlling infection, including the immune response, metabolite availability, and gut microbiome structure (reviewed in [13, 14]). Recently, a growing number of studies show an association between T2DM and infection the opportunistic pathogen *Klebsiella pneumoniae* (Kp) [15-18], yet the mechanistic connection between these diseases remains unexplored.

Kp is a frequent gut colonizer, and Kp-colonized individuals are at high risk for subsequent infection, implicating the gut as the primary reservoir for infectious Kp [19-21]. Moreover, specific gut microbial community structures are strongly associated with infection in colonized patients, and Kp gut dominance is highly associated with disease [22, 23]. It is unknown if T2DM-associated changes in the gut microbiome impact Kp colonization. Upon escape from the gut, Kp must overcome a hostile immune system and nutrient limitation to establish infection. Studies have attributed the increased susceptibility of individuals with T2DM to bacterial infection to lung glucose [24]. We have shown that amino acids are necessary for Kp lung infection [25]; however, it is unknown if this increases susceptibility to Kp infection in those with T2DM. The relative contribution of an impaired immune response, increased metabolite availability, and altered gut microbiome structure to the association between T2DM and Kp is understudied. Here, we experimentally interrogate the association between T2DM and Kp infection.

## Results

### T2DM exacerbates Kp lung infection

To measure the impact of T2DM on Kp infection, we used the BKS.Cg-*Dock7*^*m*^ +/+ *Lepr*^*db/db*^/J mice (“db/db” mice, [26, 27]), a well-characterized line of C57BLKS/J mice that phenocopy T2DM via development of chronic hyperglycemia, pancreatic β-cell atrophy and hypoinsulinemia. Both heterozygous mice (“db/+” mice) and WT C57Bl6/J mice (“+/+” mice) were used as controls for infection studies. The lung is a common primary site for Kp infection, which can lead to dissemination to the bloodstream, resulting in bacteremia and sepsis (reviewed in [28]). The hypervirulent Kp strain KPPR1 was used for lung infection experiments, as this strain is frequently used in animal infection studies and has a wide range of available genetic tools [29]. Mice were retropharyngeally inoculated with a low (10^4^ colony-forming units) or high (10^6^ colony-forming units) dose of KPPR1 to model pneumonia. Higher bacterial burdens were detected in the lungs of db/db mice compared to +/+ controls in both dose experiments (Figure 1A). db/+ mice demonstrated an intermediate phenotype in the high dose model, which suggests an impact of the reported increase in metabolic efficiency in these mice compared to +/+ controls [30]. Minimal dissemination from the lung to the liver and blood was detected across groups (Figure 1B-C).

**Figure 1.**
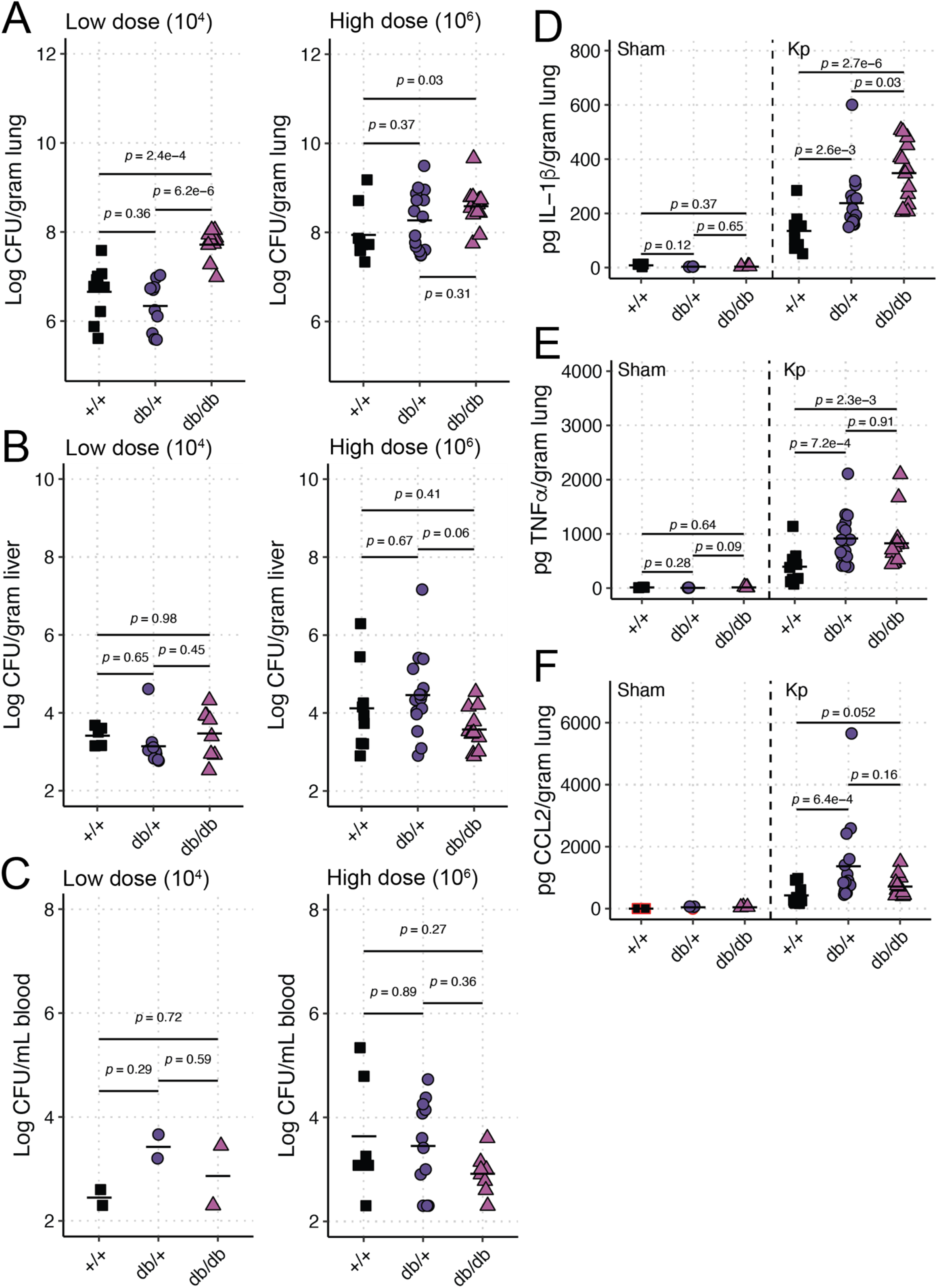
T2DM exacerbates Kp lung infection. BKS.Cg-*Dock7*^*m*^ +/+ *Lepr*^*db/db*^/J (“db/db”), BKS.Cg-*Dock7*^*m*^ -/+ *Lepr*^*db/+*^/J (“db/+”) and WT C57Bl6/J (“+/+”) mice were retropharyngeally inoculated with a low (10^4^) or high (10^6^) CFU dose of KPPR1 (n = 9-15 per group per dose). Bacterial burden in the lung (**A**), liver (**B**), and blood (**C**) was measured after 24 hours. IL-1β (**D**), TNFα (**E**), and CCL2 (**F**) were also measured 24 hours post-infection in undiluted lung homogenate of mice inoculated with 10^6^ CFU of KPPR1 by Luminex (n = 9-15 per group) or an equal volume of sterile phosphate-buffered saline (“Sham,” n = 3 per group). Each datapoint represents a single animal.

Next, we measured immune response in mice inoculated with the higher dose of KPPR1 with a Luminex assay on lung homogenate to detect cytokine production relevant to Kp infections. IL-1β (Figure 1D), TNFα (Figure 1E), and CCL2 (Figure 1F) were all significantly or nearly significantly (CCL2 *p*-value = 0.052) elevated in the lungs of db/db and db/+ mice compared to +/+ controls. IL-1β was also elevated in db/db mice compared to db/+ lungs (Figure 1D). TNFα and CCL2 levels were similar between db/db and db/+ mice (Figure 1E-F). IFNγ, IL-6, IL-10, and IL-17A were all elevated due to Kp infection, but no differences were observed between different genotypes (Figure S1). These results indicate that mice that phenocopy T2DM have more severe lung infections than healthy controls.

Metabolic diseases, including T2DM, are associated with high vascular permeability due to increases in vascular endothelial growth factor (VEGF, [31]), which may lead to increases in metabolite availability in the diabetic lung. To test if T2DM impacts lung metabolite availability via increased VEGF in our model, we first measured VEGF in lung homogenate. VEGF was significantly higher in db/db mouse lungs compared to controls at both baseline and during KPPR1 high-dose infection (Figure 2A). We then measured amino acid concentrations in bronchoalveolar lavage fluid (BALF) of uninfected mice. db/db mice exhibited a BALF amino acid composition distinct from +/+ mice (PERMANOVA *p*-value = 0.03, 1,000 permutations, Figure 2B). The db/db BALF amino acid composition did not significantly differ from db/+ mice (PERMANOVA *p*-value = 0.39, 1,000 permutations), but the BALF amino acid composition of db/+ mice marginally differed from +/+ mice (PERMANOVA *p*-value = 0.06, 1,000 permutations). The amino acids most explanatory of composition differences were taurine, creatine, glutamate, glutamine, and lysine, all of which were enriched in the db/db or db/+ mice compared to +/+ mice (Figure 2B, red arrows). db/db and db/+ mice exhibited higher concentrations of explanatory BALF amino acids compared to +/+ mice (Figure S2).

**Figure 2.**
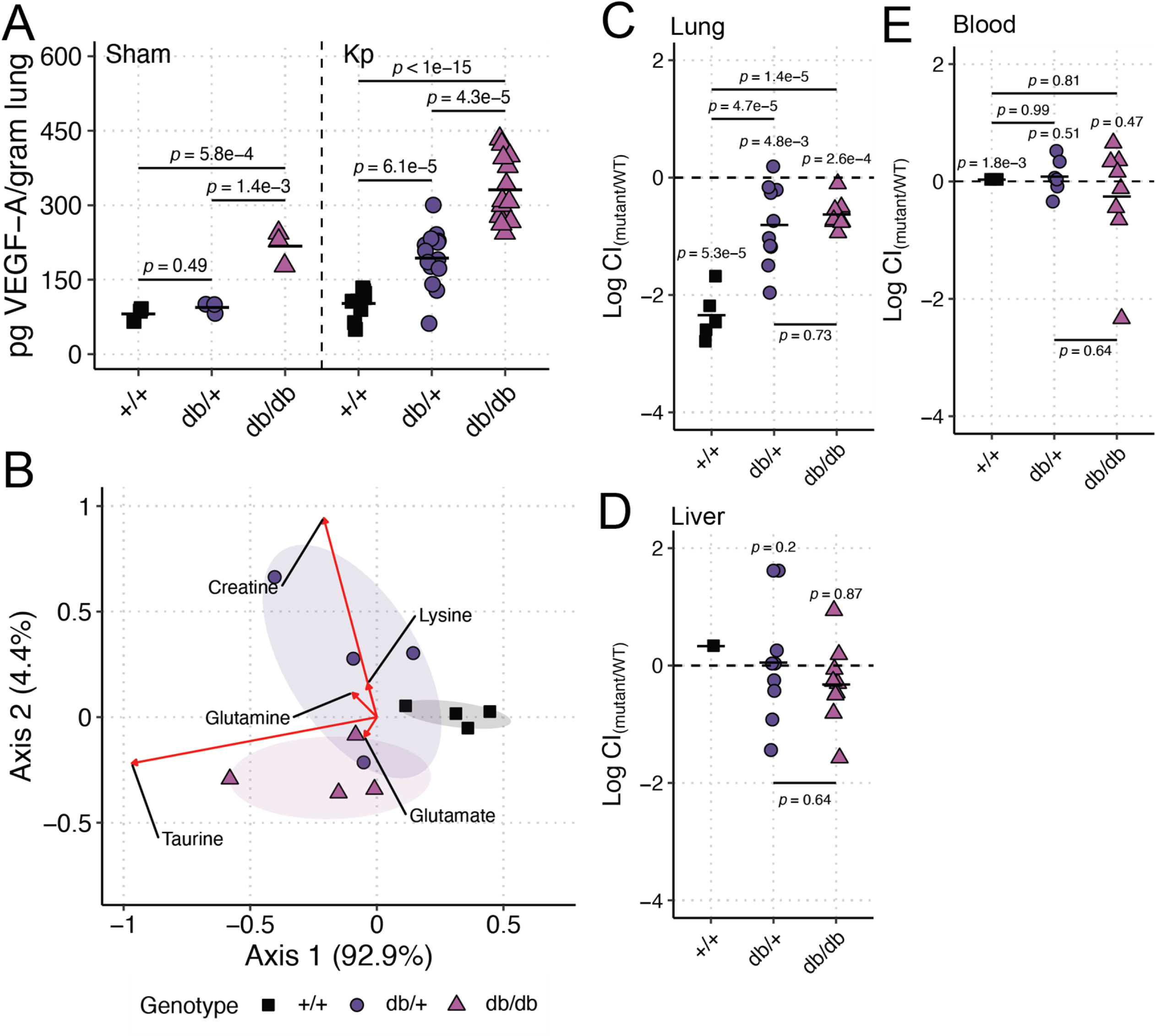
Lung amino acids are elevated in the T2DM lung in a VEGF-dependent manner, which reduces the need for metabolic flexibility during Kp lung infection. VEGF-A (**A**) was measured 24 hours post-infection in lung homogenate of BKS.Cg-*Dock7*^*m*^ +/+ *Lepr*^*db/db*^/J mice (“db/db”), BKS.Cg-*Dock7*^*m*^ -/+ *Lepr*^*db/+*^/J (“db/+”) and WT C57Bl6/J mice (“+/+”) mice inoculated with 10^6^ CFU of KPPR1 (“Kp,” n = 9-15 per group) or an equal volume of sterile phosphate-buffered saline (“Sham,” n = 3 per group). Bronchoalveolar lavage fluid was obtained from db/db, db/+, and +/+ mice (n = 4 per group), and amino acid concentrations were measured (**B**). Red arrows indicate the most explanatory amino acids differentiating the three genotypes, and the arrow direction indicates the directional impact of those amino acids. db/db, db/+, and +/+ mice were retropharyngeally inoculated a 1:1 mix of WT KPPR1 the isogenic KPPR1Δ*gltA* mutant at a dose of 10^6^ CFU (n = 5-10 per group). Bacterial burden in the lung (**C**), liver (**D**), and blood (**E**) was measured after 24 hours, and the log_10_ competitive index (Log CI) of the mutant strain compared to the WT strain was calculated for each sample. Each datapoint represents a single animal.

To confirm the bioavailability of BALF amino acids and their impact on Kp lung infection, we utilized a well-characterized Kp mutant, KPPR1Δ*gltA. gltA* encodes citrate synthase, and deletion of this gene renders KPPR1 a glutamate auxotroph [25]. We repeated our high-dose infections using a competition-based approach where KPPR1Δ*gltA* was competed against KPPR1. As expected, KPPR1Δ*gltA* was significantly less fit than its parental strain in the lung of +/+ mice (Figure 2C, Figure S3A). Interestingly, the KPPR1Δ*gltA* was significantly more fit in the lungs of db/db mice compared to +/+ mice (Figure 2C, Figure S3A), indicating higher bioavailability of glutamate family amino acids in the lungs of db/db mice. KPPR1Δ*gltA* was also significantly more fit in the lungs of db/+ mice compared to +/+ mice (Figure 2C, Figure S3A), suggesting that the db/+ mice display an intermediate phenotype as with mono-infections. As expected, GltA was dispensable in the liver and blood of all mice genotypes (Figure 2D-E, Figure S3B-C). These findings indicate that T2DM 1) exacerbates Kp lung infection and 2) increases the bioavailability of amino acids in the lung, likely through VEGF, thereby reducing the need for metabolic flexibility.

### T2DM enhances Kp gut colonization in a microbiome-dependent manner

Finally, we explored the impact of T2DM on Kp gut colonization. Using an oral gavage gut colonization model, we measured KPPR1 density in mice where the gut microbiome was either intact or disrupted by antibiotics. When the gut microbiome was intact, we observed a significant increase in Kp density in db/db mice compared to db/+ mice (Figure 3A, “No Abx”). This effect was abrogated upon administration of antibiotics (Figure 3A, “+ Abx”), indicating that elevated Kp abundance in the db/db mice is dependent on gut microbiota community structure.

**Figure 3.**
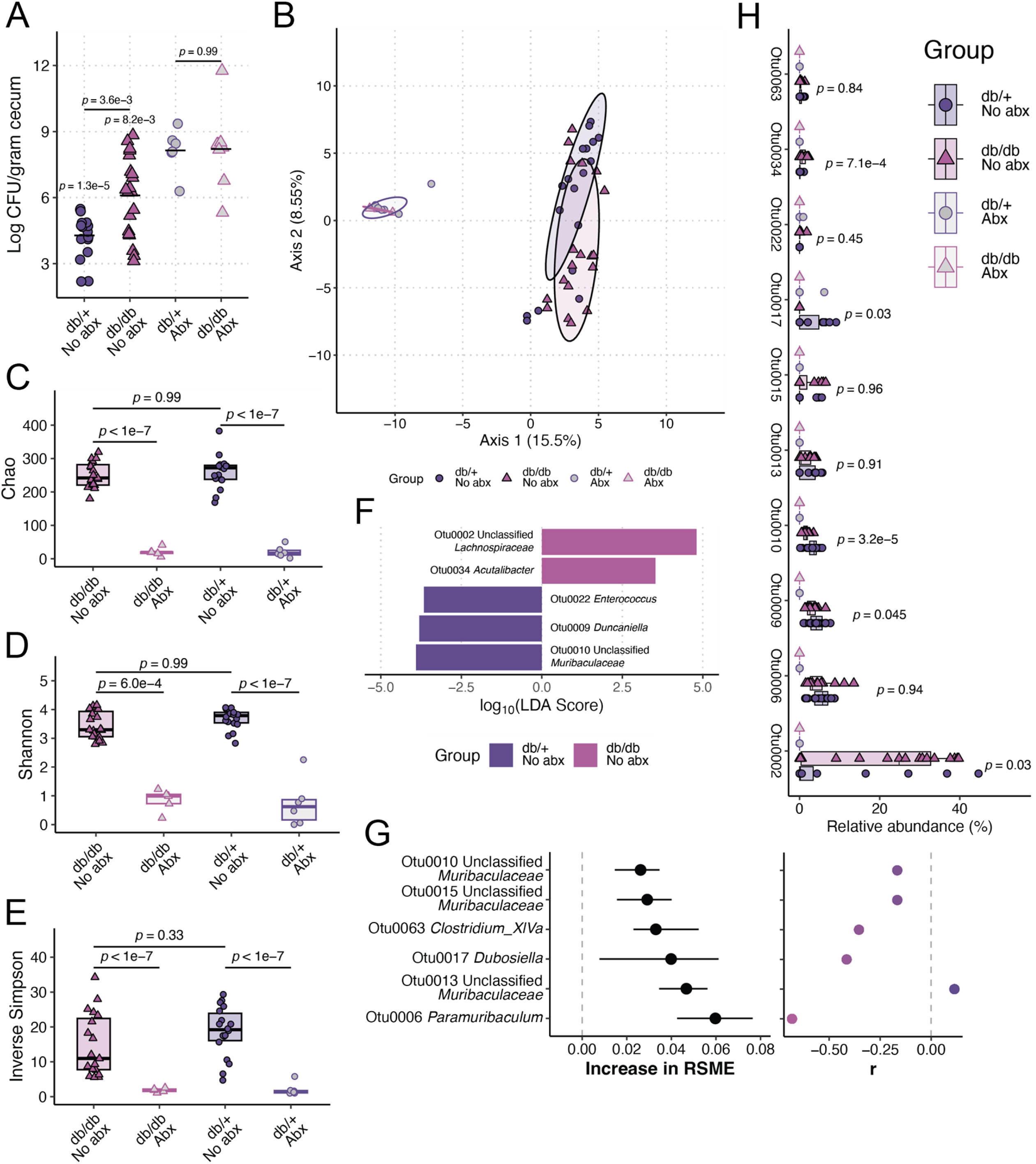
T2DM enhances Kp gut colonization in a microbiome-dependent manner. BKS.Cg-*Dock7*^*m*^ +/+ *Lepr*^*db/db*^/J mice (“db/db”) and BKS.Cg-*Dock7*^*m*^ -/+ *Lepr*^*db/+*^/J (“db/+”) were orally inoculated with 10^8^ CFU of KPPR1 with (n = 6-8 per group) and without (n = 17-20 per group) pre-administration of 0.5 g/L ampicillin via drinking water for four days. Cecal bacterial burden was enumerated 48 hours post-inoculation (**A**). Pre-inoculation fecal pellets were subjected to 16S rRNA gene sequencing to determine differences in fecal microbiome community structure. Robust Aitchison distance was used to assess the difference in β-diversity between groups (**B**). Analysis of the (**C**) Chao, (**D**) Shannon, and (**E**) Inverse Simpson α-diversity indices between groups. Linear discriminant analysis (LDA) effect size was used to identify differentially abundant (absolute log_10_(LDA score) > 3.5, *p-*value < 0.05) between db/db and db/+ mice without antibiotic administration (**F**). Regularized logistic regression and Spearman correlation were used to determine top features correlated with the cecal Kp loads of db/db and db/+ mice without antibiotic administration using OTUs as input data (**G**). Root-mean-square deviation was determined by iterating a regularized logistic regression model 100 times using 100 test data sets consisting of a random subset of samples (80%) to predict genotype status. Circles on the left graph indicate median increase in model root-mean-square deviation importance upon inclusion of each OTU in the model, and bars represent the interquartile range from 100 model iterations. Relative abundances of OTUs that were differentially abundant between db/db and db/+ fecal microbiomes or highly important features for classification of genotype status using regularized logistic regression were summarized (**H**). For panels A-E and H, each datapoint represents a single animal. For panel H, *p*-value indicates the Tukey post-hoc comparison between “db/+ No abx” and “db/db No abx” groups.

16S rRNA gene sequencing confirmed differences in fecal microbiota community structure between the intact gut microbiome of db/db and db/+ mice (PERMANOVA *p*-value = 9.99 x 10^−4^, 1,000 permutations) prior to inoculation. These differences were abrogated upon antibiotic treatment (Figure 3B). To ensure the observed differences in fecal microbiota community were not a function of the barrier of origin (the facility from which the mouse was sourced), which can impact gut microbiota community structure [32], we sourced mice from two different barriers: em08 and mp16. Genotype-dependent differences in fecal microbiota community structure between the intact gut microbiome of db/db and db/+ mice were observed regardless of the barrier of origin (Figure S4).

db/db and db/+ mice did not exhibit significant differences in α-diversity in the absence of antibiotics (Figures 3C-E), indicating that the differences between db/db and db/+ mice likely associate with dissimilarities in the relative abundance of fecal microbes and gut Kp density. Linear discriminant analysis effect size revealed differentially abundant (absolute log_10_(LDA score) > 3.5, *p-*value < 0.05) taxa between db/db and db/+ fecal microbiomes (Figure 3F). Our results were insensitive to the use of operational taxonomical units binned at 97% similarity (OTUs, Figure 3A-F) over unique amplicon sequence variants (ASVs, Figure S5).

Finally, we used supervised machine learning to measure the correlation between gut Kp density and OTU relative abundance. The median r^2^ of our models was 0.37 (interquartile range: 0.17-0.52). We determined that several OTUs were explanatory for the correlation between gut Kp density and OTU relative abundance (Figure 3G, left). Most of these correlations were negative (Figure 3G, right), suggesting a potential antagonistic relationship between Kp. Interrogation of the relative abundance of OTUs was determined to be important (Figures 3F-G) as some of these OTUs were significantly differentially abundant between db/db and db/+ mice and all were sensitive to antibiotics (Figure 3H, “Otu0002,” “Otu0009,” “Otu0010,” “Otu0017,” “Otu0035”) supporting their role in the T2DM-mediated enhancement of Kp gut colonization in a microbiome-dependent manner. Interestingly, most of the OTUs that were correlated with Kp gut density were not significantly differentially abundant between db/db and db/+ mice (Figure 3H, “Otu0006,” “Otu00013,” “Otu0010,” “Otu0015,” “Otu0063”), suggesting the effects of these bacteria that correlate with Kp gut density are independent of genotype.

Together, these data demonstrate that T2DM-associated changes to gut microbiota community structure are associated with an enhanced ability of Kp to colonize the gut when the gut microbiota community structure is unperturbed.

## Discussion

Our pneumonia and gut colonization models demonstrate that T2DM exacerbates Kp pathogenesis. Increased metabolite availability and altered gut microbiome structure may mechanistically link T2DM and Kp infection, whereas cytokine production does not. It may be the case that T2DM reduces the efficacy of immune cell killing of Kp; however, that was not tested here. Our data indicates that individuals with T2DM may be at higher risk of Kp infection due to increased susceptibility to colonization because of changes to the gut microbiota community structure. Additionally, it may be that individuals with T2DM are more susceptible than non-diabetic individuals to lung infection by Kp strains that are less metabolically flexible or efficient given the increased availability of lung metabolites. This study lays the groundwork for additional mechanistic studies to explore the connection between T2DM and the risk of infection with Kp. Future studies should aim to test these suppositions in humans and determine if glycemic control impacts Kp pathogenesis.

## Materials and Methods

### Materials, methods, and experimental protocols

A detailed description of the materials, methods, and experimental protocols for the study can be found in the Supplementary Information.

### Statistical analysis

All animal studies were replicated at least twice. Competitive indices were log_10_ transformed, and a one-sample *t-*test was used to determine significant differences from a hypothetical value of 0. All colony-forming unit (CFU) values were log_10_ transformed for analysis. A *p*-value of less than 0.05 was considered statistically significant for the above experiments. ANOVA followed by Tukey’s multiple comparisons post-hoc test was used to determine significant differences between groups. Analysis was performed using RStudio 2021.09.0+351 “Ghost Orchid” release for macOS with R v. 4.4.0.

## Data availability

The sequencing data generated in this study have been deposited in the Sequence Read Archive (SRA) database under accession PRJNA1117449 (embargoed until 01/01/2025). All other source data and code are available at https://github.com/jayvorn/Type-2-diabetes-mellitus-enhances-Klebsiella-pneumoniae-pathogenesis.

## Acknowledgements

This research was supported by work performed by The University of Michigan Microbiome Core and the Indiana University School of Medicine Biomedical Genetics Laboratory.

## Funding

This work was supported by funding from NIH/NIDDK P30 DK097512 to JV as a pilot grant from the Indiana University Center for Diabetes & Metabolic Diseases (CDMD). The funders had no role in study design, data collection and analysis, publication decision, or manuscript preparation.

## Contributions

Conceptualization: JV Methodology: KT, NTL, JV Investigation: KT, JB, KG Visualization: JV Funding acquisition: JV Project administration: JV Supervision: NTL, JV Writing – original draft: JV Writing – review & editing: KT, JB, KG, NTL, JV

## Competing interests

All authors declare that they have no competing interests.

## Notes

### Competing Interest Statement

The authors have declared no competing interest.

